# A note on noise suppression in cell-size control

**DOI:** 10.1101/098640

**Authors:** Abhyudai Singh

## Abstract

Diverse cell types employ mechanisms to maintain size homeostasis and minimize aberrant fluctuations in cell size. It is well known that exponential cellular growth can drive unbounded intercellular variations in cell size, if the timing of cell division is size independent. Hence coupling of division timing to size is an essential feature of size control. We formulate a stochastic model, where exponential cellular growth is coupled with random cell division events, and the rate at which division events occur increases as a power function of cell size. Interestingly, in spite of nonlinearities in the stochastic dynamical model, statistical moments of the newborn cell size can be determined in closed form, providing fundamental limits to suppression of size fluctuations. In particular, formulas reveal that the magnitude of fluctuations in the newborn size is determined by the inverse of the size exponent in the division rate, and this relationship is independent of other model parameters, such as the growth rate. We further expand these results to consider randomness in the partitioning of mother cell size among daughters at the time of division. The sensitivity of newborn size fluctuations to partitioning noise is found to monotonically decrease, and approach a non-zero value, with increasing size exponent in the division rate. Finally, we discuss how our analytical results provide limits on noise control in commonly used models for cell size regulation.

## Introduction

How cells regulate the timing of division to ensure *size homeostasis* and avoid getting abnormally large (or small) remains as one of the unsolved problems in biology [1]. Perhaps the simplest model of cell size regulation is one where the size of an individual cell grows exponentially over time, random cell division events occur at discrete times that partition size between two or more daughters cells. If the division timing is governed by a size-independent stochastic process (such as, a timer measuring the time elapsed since the last division event), then the statistical fluctuations in cell size grow unboundedly over time [2]. As a consequence, size homeostasis requires division timing to be regulated via a size-dependent process. Not surprisingly, uncovering mechanisms mediating size sensing and control remains a vigorous area of theoretical and experimental research in diverse organisms ranging from prokaryotes to microbial eukaryotes to plant and animal cells [3-27].

We formulate a simple stochastic model for size control, where the size grows exponentially over time throughout the cell cycle. Size dynamics is interspersed with random division events that divide size by approximately half (assuming two daughters). A key feature of the model is that the probability of division event increases as a power function of size. While such nonlinear stochastic dynamical systems are typically analytically intractable, we show that moments of the newborn size (i.e., size just after a division event) can be quantified in closed form. These results provide novel insights into how statistical variations in the newborn size are controlled by the division rate, and the randomness in size partitioning between daughters. Finally, we show how combining our analytical results with size distribution measurements allows inference of division rates.

## Stochastic formulation of cell-size control

Let *v*(*t*) denote the size of an individual cell at time *t*. Depending on the cell type, size can be quantified via different metrics, such as, cell length in rod-shaped bacteria, or cell volume/mass in animal cells. The size grows exponentially over time as

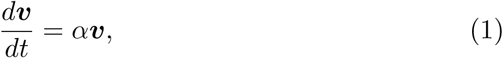

where *α* > 0 is referred to as the *exponential growth coefficient.* Cell division events occur stochastically with a *division rate f* (***v, τ***), where ***τ*** is a timer (cell-cycle clock) measuring the time elapsed from when the cell was born. More specifically, the probability of cell division occurring in the next infinitesimal time interval (*t*,*t* + *dt*] is given by *f* (***v*, *τ***)*dt*. Consistent with experimental observations [5], the division rate is a non-decreasing function in both arguments, i.e., the likelihood of division increases with size and cell-cycle progression. Whenever a cell-division event occurs, the states are reset as

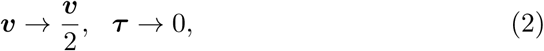

assuming that the mother cell divides into two identically-sized daughters.

It is well known that a *size-independent* division rate (*f* only depends on timer ***τ***) that yields a finite non-zero average cell size, leads to unbounded intercellular size variations [2, 28], i.e.,

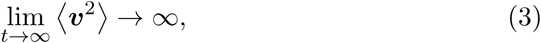

where the symbol 〈 〉 is used to denote the expected value of a random variable. Hence, a *size-dependent* division rate is an essential feature for size homeostasis. We particularly restrict ourselves to a special class of division rates that take the form of power functions

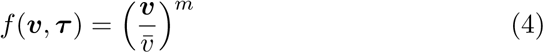

where *v̅* and m are positive constants. In the limit *m* → ∞, (4) corresponds to the well-known sizer paradigm, where the mother cell divides upon reaching a critical size threshold *v̅* [29-34]. Given that sizer implementation is never perfect (*m* is always finite), how does the extent of random fluctuation in cell size scale with the exponent *m*? Does this scaling depend on the exponential growth coefficient *α* and *v̅*? How does the scaling change if the two daughters are not identically-sized?

One approach to address these questions is to compute the statistical moments of *v*, and investigate them as a function of model parameters. For the stochastic dynamical system (1)-(4), the following set of differential equations describes the time evolution of moments

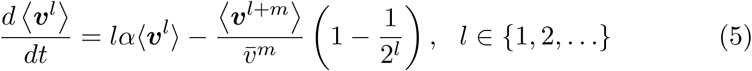

[35, 36]. Note that for any *m* > 0, (5) suffers from the problem of unclosed moment dynamics: the time evolution of any lower order moment depends on higher order moments. Hence, moments cannot be computed from (5), and approximate closure schemes are typically employed to solve such systems [35, 37-48].

## Statistical moments of newborn cell size

Note that considerable cell-to-cell variations in *v* can simply be attributed to the fact that different cells are in different stages of their cell cycle. Taking this into account, perhaps a more meaningful quantity is the newborn cell size (*v* conditioned on *τ* = 0). We reformulate the size control model presented in the previous section into a discrete-time stochastic system that tracks the newborn size across generations

Consider a single cell undergoing cycles of growth and division. Let random variable ***V_i_*** denote the newborn size during the start of the *i^th^* cell cycle. Then, given that size increases exponentially over time, and the division rate *f* (*v*, *τ*), the time *T_i_* from the start of the *i^th^* cell cycle when division is triggered follows the probability density function (pdf)

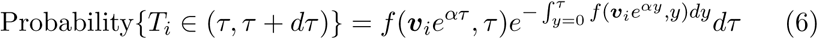

[2, 49, 50]. Note that a constant function *f* in (46) would correspond to exponentially distributed *T_i_*. In our case, *f* takes the following time-varying form based on (4)

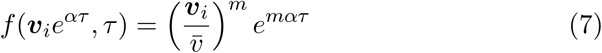

that results in *T_i_* with lower statistical variations than an exponential random variable (i.e., memory in division timing), and dependent on *v_i_* (i.e., larger newborns divide earlier as compared to smaller newborns). Having determined the time to division, the newborn size in the next cycle is

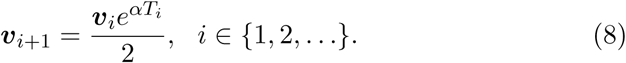

This discrete-time stochastic system yields the following *l^th^* order moment of ***v**_i+1_* conditioned on ***v**_i_*

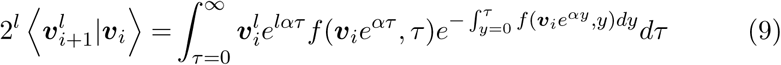

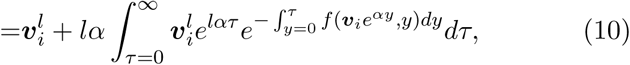

where the first integral is simplified via integration by parts. Next, we show how this recurrence equation can be solved in moments space.

### Linear dependence of division rate on cell size

We first restrict ourselves to *m* = 1 in the division rate (4), which corresponds to the probability of division-event occurrence increasing linearly with size. In this case, using

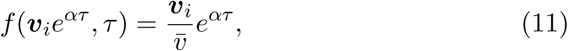

(10) can be written as

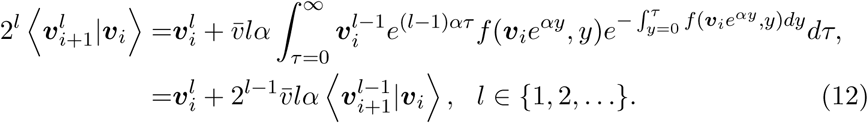

Now unconditioning on *V_i_* and taking the limit *i*→∞

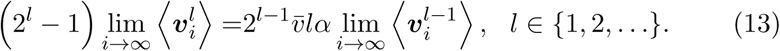

The average newborn size at steady state is obtained by substituting *l* = 1

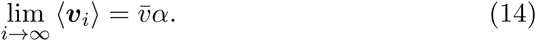

In general, (13) can be solved iteratively to yield

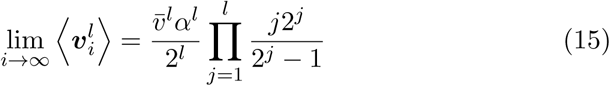

*Thus, in spite of unclosed moment dynamics, the moments of newborn size can be computed exactly in closed form*.

One observation from (15) is that if we define a normalized size (size scaled by its average)

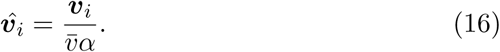

then all moments of ***v**̂_i_* become invariant of *α* and the mean cell size. This was dramatically seen in experiments with *E. coli*, where the newborn size distributions across different growth conditions collapse on top of each other once scaled by their respective means [51]. The noise in newborn size, as quantified by its Coefficient of Variation (*CV*) squared, is give by

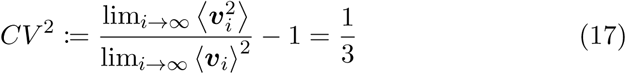

and is completely independent of model parameters *v̅* and *α*. *CV*^2^ computed in (17) implies a standard deviation that is ≈ 58% of the average size. An interesting point to note is

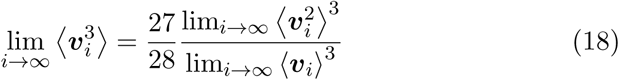

Since for any lognormally distributed random variable *Y*,

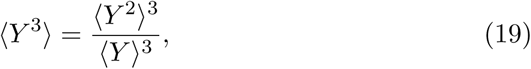

(18) suggests that as far as the third-order moment is concerned (i.e, skewness), the newborn size at equilibrium is approximately lognormal. Next, we consider the general case of any exponent *m* in (4).

### Division rate as a power function of cell size

Before considering the general case it is worthwhile to note that as *m* → ∞, the size just before, and after division, approaches *v̅* and *v̅*/2 with probability one, respectively. Thus, one expects the noise in newborn size *CV*^2^ → 0 as *m* → ∞, but the question remains as to how fast it goes to zero.

Our strategy in this case relies on making the division rate linear through the following transformations

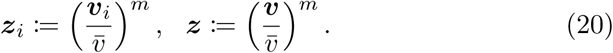

Given that *v(t)* increases exponentially between division events, *z(t)* follows similar dynamics, but with a different exponential growth coefficient

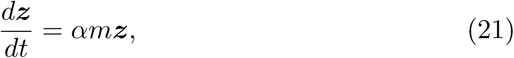

with resets

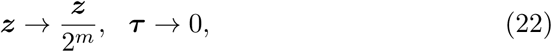

that are activated at the time of division. This transformation leads to the following discrete-time system

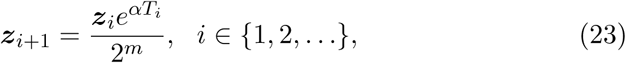

where *T_i_* is given by (46) with division rate *f* = *z*. Following steps similar to the previous section yield the following moments for *z_i_*

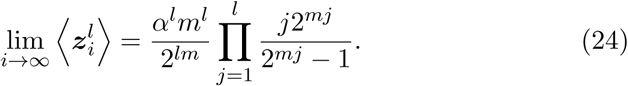

### The case of exponent *m* ≤ 1

To obtain the moments of the newborn size using (24) we first consider the case *m* = 1/*n* ≤ 1, where *n* is a positive integer. In this case,

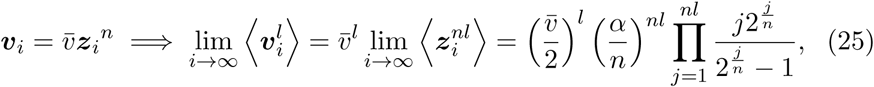

which yields the following noise in newborn size

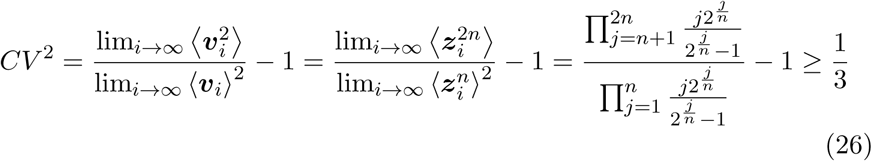

As expected from (18), *CV*^2^ = 1/3 for *m* = 1 (*n* = 1). Moreover, *CV*^2^ increases with decreasing *m*, and *CV*^2^ → ∞ as *m* → 0 (*n* → ∞). This latter result is consistent with the fact that a size-independent division rate yields infinite variance in newborn size [2, 52].

### The case of exponent *m* ≥ 1

Experimental quantification of size distributions in both *E. coli* and mammalian cells report *CV*^2^ values considerably smaller than 1/3 [5, 51, 53-55]. Since *CV*^2^ ≥ 1/3 for *m* ≤ 1, and *CV*^2^ → 0 as *m* → ∞, suggests that the physiological values of *m* are much larger than one. When *m* ≥ 1

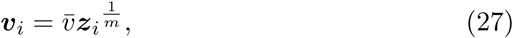

where all steady-state moments of *z_i_* are given by (24). While exact analytical formulas for the moments of *v_i_* are not available in this case, we show how simple approximate formulas can be derived.

The first approximation relies on Taylor series. Assuming fluctuations in *z_i_* are small, expanding the right-hand-side of (27) around the steady-state average lim*_i_*_→∞_ 〈*z^i^*〉

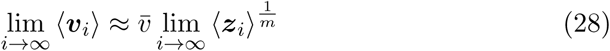

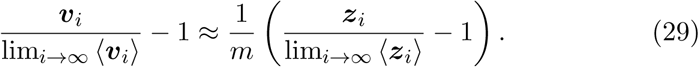

Squaring both sides of (29) and taking the expected value yields

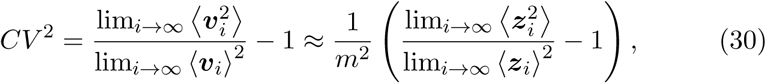

which using (24) yields the following approximation

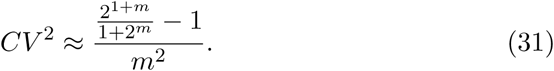

While this approximation is exact for *m* = 1 (*CV*^2^ = 1/3), it predicts noise in newborn size to monotonically decrease with increasing *m* with the asymptote

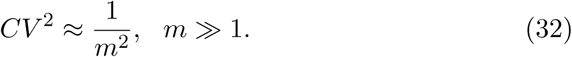

The second approximation works for large values of *m*. Assuming *m* ≫ 1, then from (20) and (24)

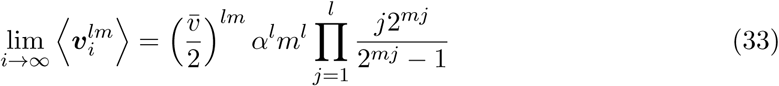

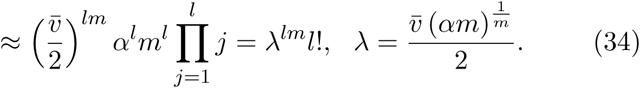

The moments in (34) are consistent with the newborn size at equilibrium following a Weibull distribution with scale parameter *λ* and shape parameter *m* [33]. For such a Weibull distribution the coefficient of variation squared is given by

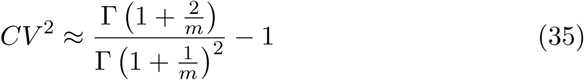

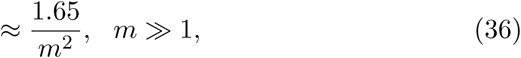

where Γ denotes the Gamma function. Both the above approximations are plotted in Fig. 1. While (31) is much more accurate for 1 ≤ *m* ≤ 5, (35) is better for *m* ≥ 6. Note that both approximations imply a 1/*m* scaling of the newborn size *CV*.

**Figure 1:**
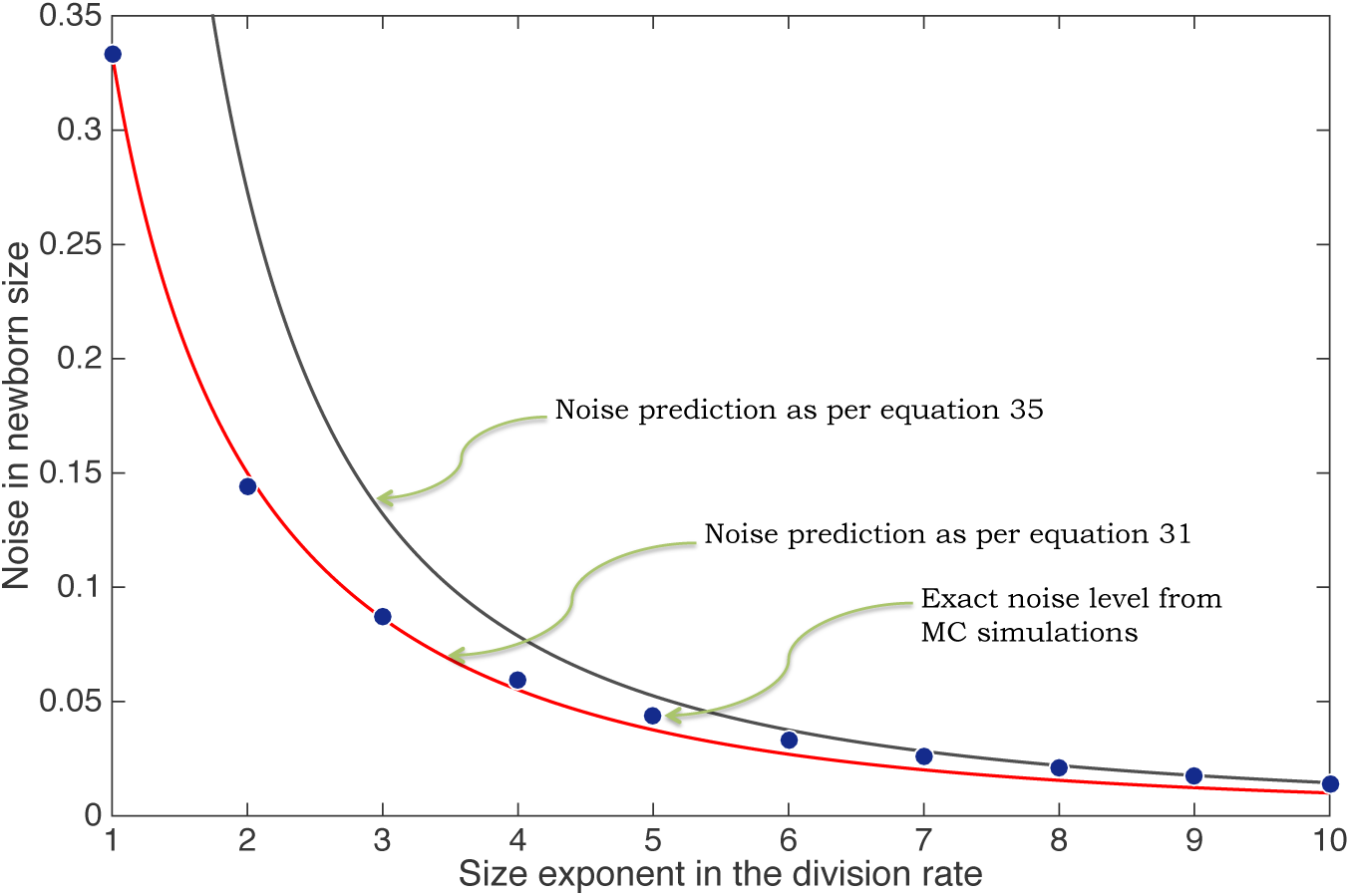
Noise in newborn cell size, as qauntified by its steady-state coefficient. of variation squared, is plotted as function of the size exponent *m* in the division rate (4). For each *m*, the exact noise level is determined by running 10^7^ Monte Carlo simulations of the discrete-time system (23). While the approximation (31) is more accurate for small *m*, approximation (35) performs better for large *m*.

## Incorporating partitioning errors

Our analysis up till now has assumed perfect symmetric division of a parent cell into two daughters. Next we consider more realistic partitioning scenarios that allow for stochastic differences between daughter cell size. In particular, at the end of *i^th^* cell cycle the size is reset as

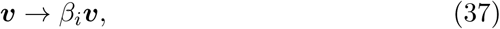

where *β_i_* are independent and identically distributed random variable drawn from a size-independent beta distribution with mean 〈*β_i_*〉 = 1/2. Noise in the partitioning process is quantified via the coefficient of variation squared of *β_i_*, 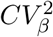. Experimentally, 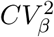 is quantified by tracking sizes of both daughters via

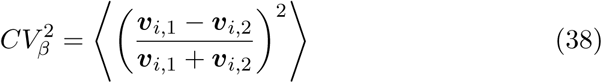

where *v_i_*,_1_ and *v_i_*,_2_ denote the two daughter cell sizes just after division. The value of 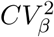 is estimated to be 10^−2^ for mammalian cells [5], and 10^−3^ — 10^−4^ for *E. coli* [53]. The low partitioning noise in *E. coli* is due to the precise positioning of the septal ring at the cell midpoint using the Min protein system [56, 57].

In the limit *m* → ∞, i.e., division occurs upon attainment of a critical size *v̅*, the newborn size is given by

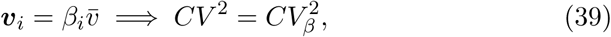

and hence, newborn size fluctuations are completely determined by the randomness in the partitioning process. Does *CV*^2^ become more or less sensitive to 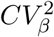 for lower values of *m*? To address this question, we consider a linear division rate (*m* = 1), where moments can be determined exactly. With inclusion of random partitioning, the stochastic dynamics of the newborn size is described by the discrete-time system

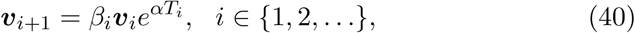

which yields the following moments of *v_i_*_+1_ conditioned on *v_i_*

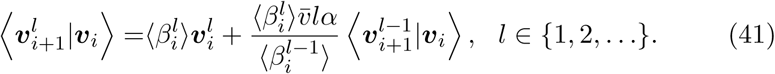

Unconditioning on *v_i_* and taking the limit *i* → ∞

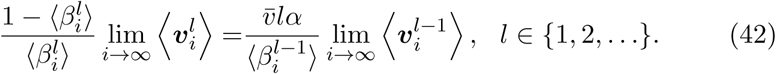

Solving this recurrence equation leads to the following moments

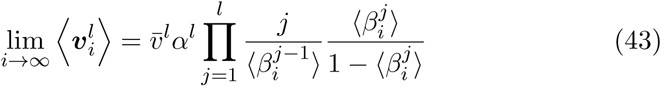

Finally, substituting *l* = 1, 2 and using 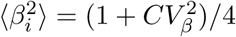 results in

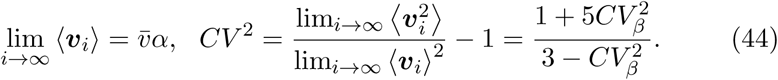

Our results show that for *m* = 1, *CV*^2^ increases monotonically with 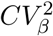, but unlike (40) the dependence is nonlinear. For small values of 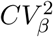, *CV*^2^ can be decomposed as

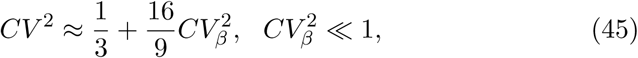

where the terms on the right represent noise contributions from random division events and stochastic partitioning, respectively. This result implies that the sensitivity of newborn size fluctuations to random partitioning is 16/9 ≉ 1.78 for *m* = 1, and higher than the sensitivity for *m* = ∞ (40).

## Conclusion

In summary, we have analyzed a stochastic model for size control, where the likelihood of division events takes the form of power functions. Exact and approximate formulas quantifying the magnitude of fluctuations in the newborn size were derived and reveal the following key insights:

- In the absence of partitioning noise 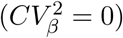, the extent of newborn size fluctuations is predicted to scale inversely with the exponent *m* in the division rate.
- Newborn size follows a lognormal distribution for values of *m* close to one, and a Weibull distribution for *m* ≫ 1.
- Consistent with experimental observations [51], the normalized size defined by (16) has moments that are independent of the exponential growth coefficient and the mean cell size.
- The inclusion of random partitioning 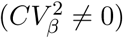 creates an additional noise term that increases nonlinearly with 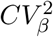. The senstivity of this noise term to 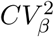 decreases with increasing *m*, and approaches a value of one for *m* → ∞.

Can the results presented here be generalized to other forms of division rates? Recent size measurements in prokaryotes have uncovered the adder strategy for size homeostasis, where division is triggered after the cell accumulates a constant size from birth [51, 53, 58-60]. In our framework, the adder can be implemented via

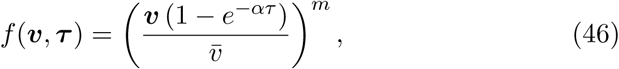

and a large enough *m* corresponds to division occurring after adding a size *v̅* to the newborn size [61]. Note that (46) has timer control in addition to size control that prevents a cell to divide just after birth (i.e., *f* (*v*, 0) = 0). This added timer control ensures that for perfect size partitioning, the noise in newborn size is always lesser than the sizer strategy (4) for the same value of *m* [61]. This result has important implication for estimation of *m* from data. For example, [51] showed that the *CV* of *E. coli* newborn was ≈ 15% across growth conditions. Using a *CV*^2^ = .023, Fig. 1 provides a value of *m* ≈ 7 for the division rate (4), and this implies a *m* < 7 for the adder strategy. With the increasing focus of measuring size at a single cell resolution, our theoretical results provide an important basis to understand origins of size fluctuations, and in turn use these noise measurements to uncover size control principles.

## Acknowledgments

This work is supported by the National Science Foundation Grant DMS-1312926.

